# Altered Mediator dynamics during heat shock in budding yeast

**DOI:** 10.1101/2020.08.25.267088

**Authors:** Debasish Sarkar, Z. Iris Zhu, Emily Paul, David Landsman, Randall H. Morse

## Abstract

The Mediator complex is central to transcription by RNA polymerase II (Pol II) in eukaryotes. In yeast, Mediator is recruited by activators via its tail module and then facilitates assembly of the pre-initiation complex (PIC), including Pol II, setting the stage for productive transcription. Mediator occupies proximal promoter regions only transiently prior to Pol II escape; interruption of the transcription cycle by inactivation or depletion of Kin28 inhibits Pol II escape and stabilizes Mediator occupancy at promoters. However, whether Mediator occupancy and dynamics differ for gene cohorts induced by stress or alternative growth conditions has not been examined on a genome-wide scale. Here we investigate Mediator occupancy following heat shock or CdCl_2_ induction, with or without depletion of Kin28. We find that Pol II occupancy exhibits similar dependence on Mediator under normal and heat shock conditions; however, Mediator occupancy does not increase upon Kin28 depletion at most genes active during heat shock, indicating altered dynamics. Furthermore, Mediator occupancy persists at genes repressed by heat shock or CdCl_2_ induction and exhibits peaks upstream of the proximal promoter whether or not Kin28 is depleted, suggesting that Mediator is recruited by activators but is unable to engage PIC components at these repressed targets. Finally, we show a reduced dependence on PIC components for Mediator occupancy at promoters after heat shock, further supporting an altered dynamics or stronger engagement with activators under these conditions.

## Introduction

Transcription of protein-coding genes in eukaryotes is a complex process involving recruitment of coactivators, assembly of the pre-initiation complex (PIC), including RNA polymerase II (Pol II), and transition from promoter melting and transcription initiation to productive transcriptional elongation (Schier and Taatjes 2020). The Mediator complex, a large, multi-subunit complex present in plants, animals, and single-celled eukaryotes, is central to this process (Jeronimo and Robert 2017; Soutourina 2018). In yeast, Mediator has been divided into head, middle, tail, and cyclin-CDK modules on the basis of structural and genetic data (Plaschka et al. 2016). Although recent, high-resolution cryo-EM investigations have revised and added nuance to Mediator structure (Tsai et al. 2014; Wang et al. 2014; Plaschka et al. 2015), it remains the case that functional attributes of yeast Mediator segregate, to some extent, to distinct modules (van de Peppel et al. 2005; Knoll et al. 2018).

Functionally, activators recruit yeast Mediator to upstream activation sites (UAS regions) via the tail module (in particular the tail module triad of Med2-Med3-Med15); subunits in the middle and head module then engage PIC components, including Pol II, to facilitate PIC assembly (Jeronimo and Robert 2017; Knoll et al. 2018; Soutourina 2018). This process entails bridging between the UAS and the proximal promoter, where the PIC is assembled, by a single Mediator complex (Jeronimo et al. 2016; Petrenko et al. 2016). Mediator ChIP signal at UAS regions is variable in strength and not well correlated with transcription levels, while under normal conditions, Mediator association with proximal promoter regions is transient and difficult to detect by ChIP (Fan et al. 2006; Fan and Struhl 2009; Jeronimo and Robert 2014; Wong et al. 2014; Paul et al. 2015b). For example, genes encoding ribosomal proteins (RP genes) depend on Mediator for Pol II recruitment under normal growth conditions, yet many exhibit weak Mediator ChIP signal in spite of being transcriptionally highly active. The conclusion that Mediator does occupy proximal promoters, if normally only transiently, is based on experiments showing that depletion or inactivation of Kin28, which inhibits promoter escape by Pol II, results in robust Mediator ChIP signal at gene promoters (Jeronimo and Robert 2014; Wong et al. 2014). Consistent with this interpretation, Mediator occupancy at promoters under conditions of Kin28 depletion is reduced by simultaneous depletion of Pol II, Taf1, or TBP; in the latter case, increased Mediator ChIP signal is observed at UAS sites, indicating that TBP is required for transit of Mediator to promoters from its initial sites of recruitment (Knoll et al. 2018).

Mediator occupancy measured by ChIP during the normal transcription cycle (i.e. with Kin28 active) is much higher at induced genes that are controlled by strong activators, such as those activated by heat shock or growth in galactose (Fan et al. 2006; Fan and Struhl 2009; Kim and Gross 2013). This could reflect stronger interactions between Mediator and activators, and hence less dependence on PIC components or other factors for stabilizing Mediator occupancy, at strongly induced genes than at constitutively active genes during growth in rich medium (YPD). Increased Mediator occupancy has been observed at a few individual promoters induced by heat shock following Kin28 depletion (Petrenko et al. 2016); however, no genome-wide analysis has been reported examining whether promoter occupancy by Mediator is stabilized by Kin28 depletion at strongly induced genes or depends on PIC components as it does in yeast grown in rich medium (YPD). In this work, we examine genome-wide Mediator occupancy following two perturbations, heat shock and CdCl_2_ administration, that induce large and rapid genome-wide transcriptional responses.

## Results

### Pol II recruitment upon heat shock depends on Mediator

To identify genes most active after heat shock and most highly induced by heat shock, we performed ChIP-seq to determine Pol II occupancy in the commonly used laboratory strain BY4741, and in YFR1321, the parent strain to that used for anchor-away of Kin28, before and after 15 and 30 min of heat shock. Heat shock resulted in increased Pol II occupancy at targets of Hsf1, the primary transcription factor responding to heat shock, and the stress responsive activators Msn2 and Msn4, while Pol II occupancy was nearly completely lost from RP genes, consistent with previous determinations of mRNA levels and nascent RNA production following heat shock (Figure 1A) (Warner 1999; Gasch et al. 2000; Causton et al. 2001; Pincus et al. 2018). We observed a bias for Pol II occupancy towards the 5’ ends of genes after heat shock, as has been noted previously (Kim et al. 2010; McKinlay et al. 2011). K-means clustering revealed two groups of genes strongly up-regulated by heat shock; these groups were both enriched for association with Hsf1 and Msn2 and Msn4 (Figure 1B). Transcription induced by Hsf1 has been reported to occur independently of Mediator (Lee and Lis 1998; McNeil et al. 1998), but we found a similar 2-3 fold reduction in Pol II occupancy at Hsf1 and Msn2/4 targets after heat shock upon depletion of the Mediator head module subunit Med17, using the anchor away method (see next section), as at other genes in the absence of heat shock, in agreement with recent results from the Struhl lab (Petrenko et al. 2017) (Figure 1C).

**Figure 1.**
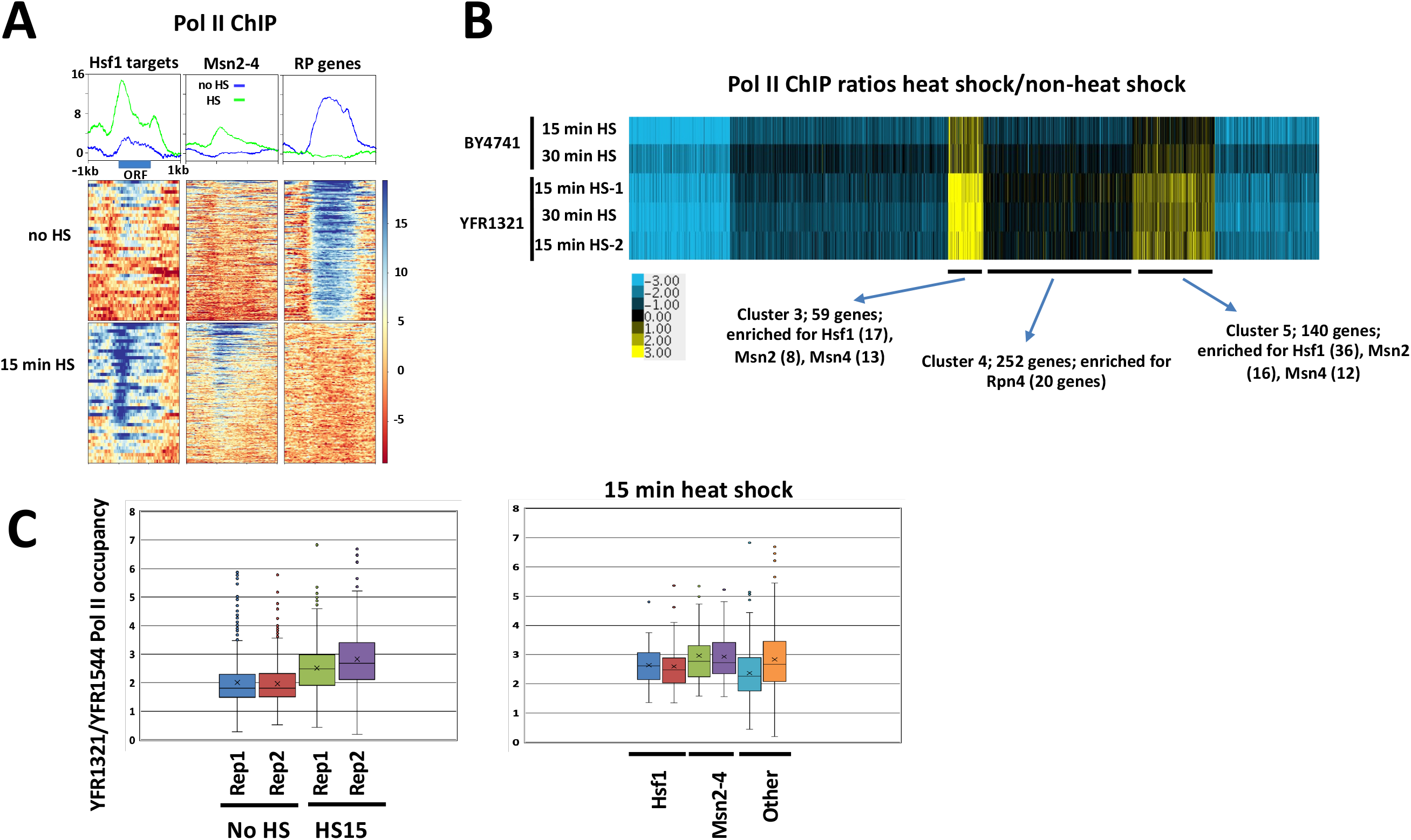
Pol II recruitment following heat shock. (A) Heat maps and line graphs depicting normalized Pol II occupancy in YFR1321 cells before and after 15 min heat shock at 42 Hsf1 targets and 213 Msn2-4 targets (see Methods) and 137 RP genes. (B) K-means clustering (K=6) was performed for the ratio of Pol II occupancy (normalized for gene length) before and after heat shock, using the 1000 genes having highest Pol II occupancy under non-heat-shock conditions plus the 300 gene having highest occupancy after heat shock (1179 ORFs, due to overlap between the two sets). Enrichment for TFs in individual clusters, as shown, derived from CERES (Morris et al. 2010). (C) Box and whisker plots depicting ratios of Pol II occupancy (normalized for gene length) with and without heat shock, as indicated, in the *med17*-AA strain (YFR1544) and parent strain (YFR1321). Left panel depicts replicate experiments examining the 886 genes with highest Pol II occupancy in the absence of heat shock (three outliers with ratios >8 in replicate 1 were removed for clarity) and 294 genes having highest occupancy after heat shock. Right panel depicts the 294 genes having highest Pol II occupancy after heat shock, divided into Hsf1 targets (34 genes), Msn2-4 targets (48 genes), and other genes (212 genes).

### Effect of Kin28 depletion on Mediator recruitment with and without heat shock

To examine Mediator dynamics at genes induced upon heat shock, we performed ChIP-seq against Med15, from the Mediator tail module triad, in yeast engineered to allow depletion of Kin28 from the cell nucleus using the anchor away method (*kin28-AA* yeast), and in the parent strain, YFR1321 (Jeronimo and Robert 2014; Wong et al. 2014; Knoll et al. 2018). (We used epitope-tagged derivatives of these strains (see Methods) to allow ChIP of Med15 and Med18; for simplicity, we refer to these strains by the parent strain names throughout the text and figure legends.) These strains harbor a *tor1-1* mutation that abrogates the stress response normally accompanying exposure to rapamycin (Haruki et al. 2008). The *kin28-AA* yeast strain expresses Kin28 with a C-terminal FRB tag and the ribosomal subunit RPL13A C-terminally tagged with the FKBP12 fragment; upon administration of rapamycin, Kin28-FRB is tightly coupled to RPL13A-FKBP12 and is evicted from the nucleus following nuclear processing of RPL13A. The parent strain is identical except for lacking the FRB tag. Yeast cultures were grown to early log phase (A_600_~0.6-0.8), treated with rapamycin for 1 hr to allow Kin28 depletion, and then subjected to heat shock by a rapid temperature shift for 15-30 minutes prior to cross-linking for ChIP-seq analysis. Previous work has shown that 1 hr of rapamycin treatment results in efficient depletion of target protein and of Kin28 in particular, and robust recruitment of Mediator has been observed after 5-30 minutes of heat shock (Haruki et al. 2008; Kim and Gross 2013; Wong et al. 2014; Knoll et al. 2018).

In the absence of heat shock, depletion of Kin28 resulted in an increase in Med15 ChIP signal at promoters of Hsf1 targets and RP genes, consistent with prior work (compare “Parent no HS” and “kin28-AAR no HS” in Figure 2A; see also Figure S1A for biological replicate experiment) (Jeronimo and Robert 2014; Wong et al. 2014). Many Hsf1 target genes are active even in the absence of heat shock (Gross et al. 1990; Solis et al. 2016; de Jonge et al. 2017; Pincus et al. 2018), so the association of Mediator with promoters of Hsf1 targets in the absence of heat shock, when Kin28 is depleted, is not unexpected. Heat shock resulted in an increased Med15 ChIP signal at Hsf1 and Msn2/4 targets in both the parent strain and the *kin28-AA* strain (Figure 2A). Quantitation of Med15 ChiP signal at RP genes and at the ~300 non-RP gene promoters having highest Med15 occupancy in non-heat-shocked cells after Kin28 depletion revealed an increase in signal in *kin28AA* compared to WT yeast, as expected; this increase was especially pronounced for RP genes (Figure 2B and Figure S1B). In contrast, increased Med15 occupancy was not observed upon Kin28 depletion in heat-shocked cells, either at Hsf1 targets or the 300 genes most highly occupied by Med15 in *kin28AA* yeast after heat shock (Figure 2A-B and Figure S1A-B). Examination of the 300 non-RP gene promoters with highest Pol II occupancy (in non-heat shocked and heat-shocked YFR1321 yeast) yielded similar results (Figure S1C), as did comparison of genes having similar Pol II occupancy in the absence and presence of heat shock (Figure S1D).

**Figure 2.**
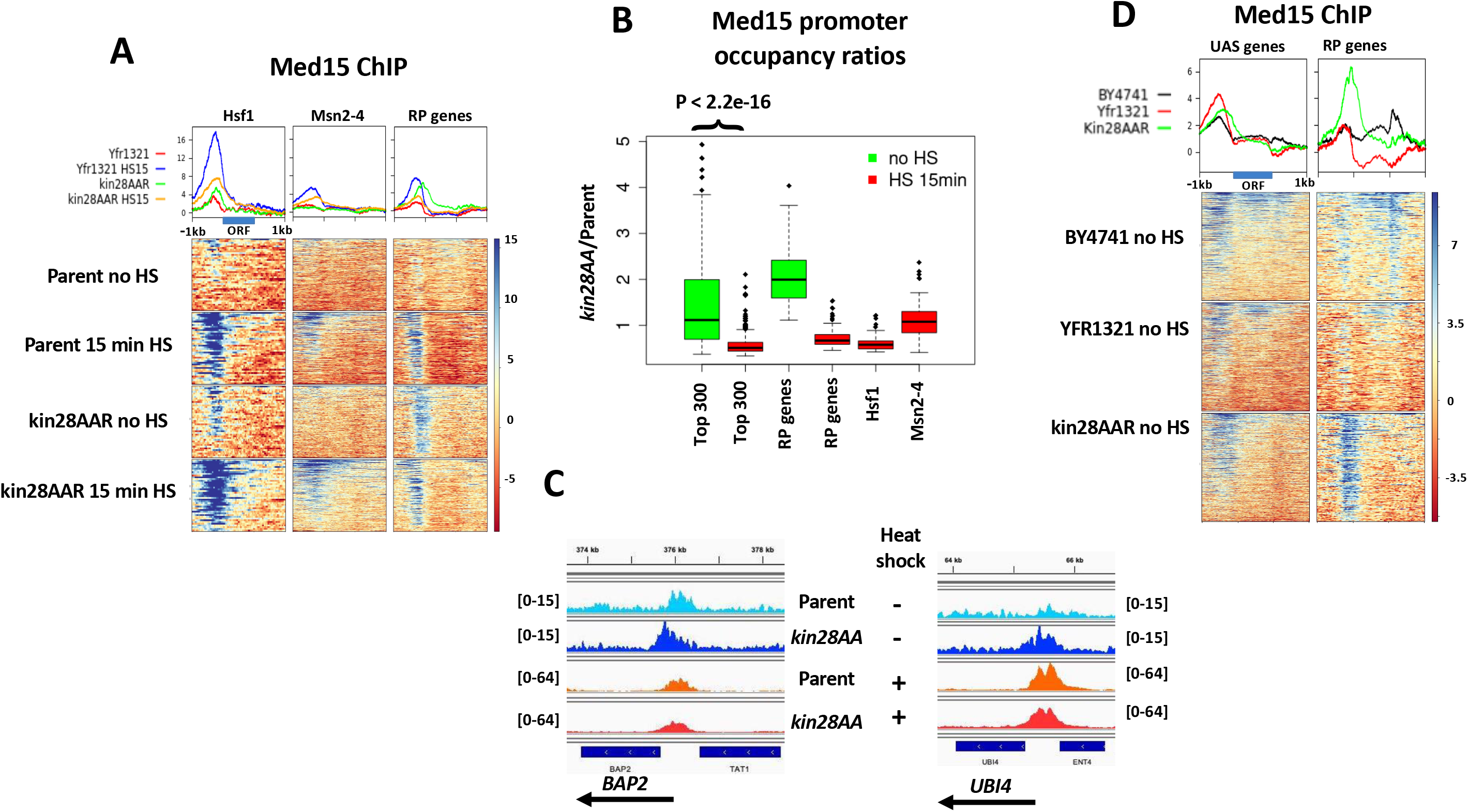
Effect of heat shock on Mediator association. (A) Heat maps and line graphs depicting normalized occupancy of the Mediator tail module subunit, Med15, in *kin28AA* yeast treated with rapamycin (“kin28AAR”) and the parent strain YFR1321, also treated with rapamycin, before and after 15 min heat shock, at 42 Hsf1 targets and 213 Msn2-4 targets (see Methods) and 137 RP genes. (B) Box and whisker plots showing the ratios of Med15 occupancy with and without Kin28 depletion for the ~300 genes showing highest Med15 occupancy in Kin28-depleted cells without or with heat shock; ratios are also shown for RP genes (137 genes), and for Hsf1 targets (42 genes) and Msn2-4 targets (213 genes) in heat-shocked cells. The p-value for comparison of the ratios for the top 300 genes with and without heat shock was calculated using the Wilcoxon rank sum test. (C) Browser scans showing Med15 occupancy upstream of *BAP2* and *UBI4* in *kin28AA* yeast and the parent strain YFR1321, both treated with rapamycin, with and without heat shock. *UBI4* is a target of Hsf1, while *BAP2* is not a target of Hsf1 or Msn2-4. Scale, in reads per million mapped reads, is indicated for each scan. (D) Heat maps and line graphs depicting occupancy of the Mediator tail module subunit, Med15, in BY4741 yeast, *kin28AA* yeast treated with rapamycin (“kin28AAR”) and the parent strain YFR1321, also treated with rapamycin, in the absence of heat shock, at 498 “UAS” genes (see text) and 137 RP genes.

Consistent with the trends evident from Figures 2A-B, stronger induction of Med15 occupancy upon Kin28 depletion under non-heat-shock than under heat-shock conditions was observed upon inspection of browser scans at many genes (Figure 2C; Figure S2, *YDJ1*). At the same time, some promoters behave counter to this trend, reflecting the range of altered Med15 occupancy observed upon Kin28 depletion (Figure 2B; Figure S2, *TPS1*). Variable effects on Med15 occupancy increase upon Kin28 depletion were observed for Hsf1 and Msn2/4 targets as well as for genes induced by heat shock that are not known to be targets of either of these stress-responsive activators (Figure S2; Table S1). Gene-specific increase in Mediator ChIP signal upon Kin28 depletion or inactivation is similarly evident in previous reports, but the cause of this remains unexplained (Jeronimo and Robert 2014; Knoll et al. 2018).

Under normal growth conditions in rich medium, many active gene promoters, most conspicuously those of many RP genes, exhibit little or no association with Mediator as measured by ChIP (Jeronimo and Robert 2014; Wong et al. 2014; Paul et al. 2015b). Some active genes, however, do exhibit Mediator ChIP signal at UAS regions under normal growth conditions; depletion or inactivation of Kin28 results in a shift in Mediator occupancy at these genes from UAS to proximal promoter (Jeronimo and Robert 2014; Wong et al. 2014). Correspondingly, we observed a shift towards the proximal promoter with little change in intensity of the Med15 ChIP signal upon Kin28 depletion in the absence of heat shock at a set of 498 genes, which we refer to as “UAS genes”, identified as exhibiting Mediator occupancy of UAS regions under normal growth conditions (Figure 2D) (Jeronimo et al. 2016). In contrast, RP genes show greatly increased Mediator occupancy upon Kin28 depletion, along with a shift in the peak closer to the TSS (Figure 2D).

Taken together, these results indicate that Hsf1 targets that are active under non-heat-shocked conditions behave similarly to the large cohort of genes, exemplified by RP genes, that exhibit little or no Mediator signal unless Kin28 is depleted, while under heat shock conditions, Hsf1 and Msn2/4 targets behave more like the “UAS genes” at which Mediator ChIP signal is seen under normal growth conditions and exhibits modest increase upon depletion of Kin28.

### Mediator remains associated with promoters of genes repressed by heat shock

We were surprised to note a prominent Med15 peak upstream of RP genes following heat shock (Figure 2A), despite the near absence of Pol II occupancy (Figure 1A). This peak was unaffected in position by Kin28 depletion in heat shocked cells, and was shifted slightly upstream relative to the Med15 peak observed following Kin28 depletion in the absence of heat shock (Figure 2A and Figure 3A). Thus, upon heat shock, Mediator association with RP genes, under conditions of depleted Kin28, shifts from a proximal promoter location to an upstream position that closely coincides with that of the RP gene activator Rap1 (Fig. 3A and see below). A similar shift upstream in *kin28AA* yeast following heat shock was seen at RP genes for the Mediator head module Med18, also known as Srb5, with the signal also showing a somewhat greater decrease in intensity compared to that for Med15 (Figure 3B; Figure S3).

**Figure 3.**
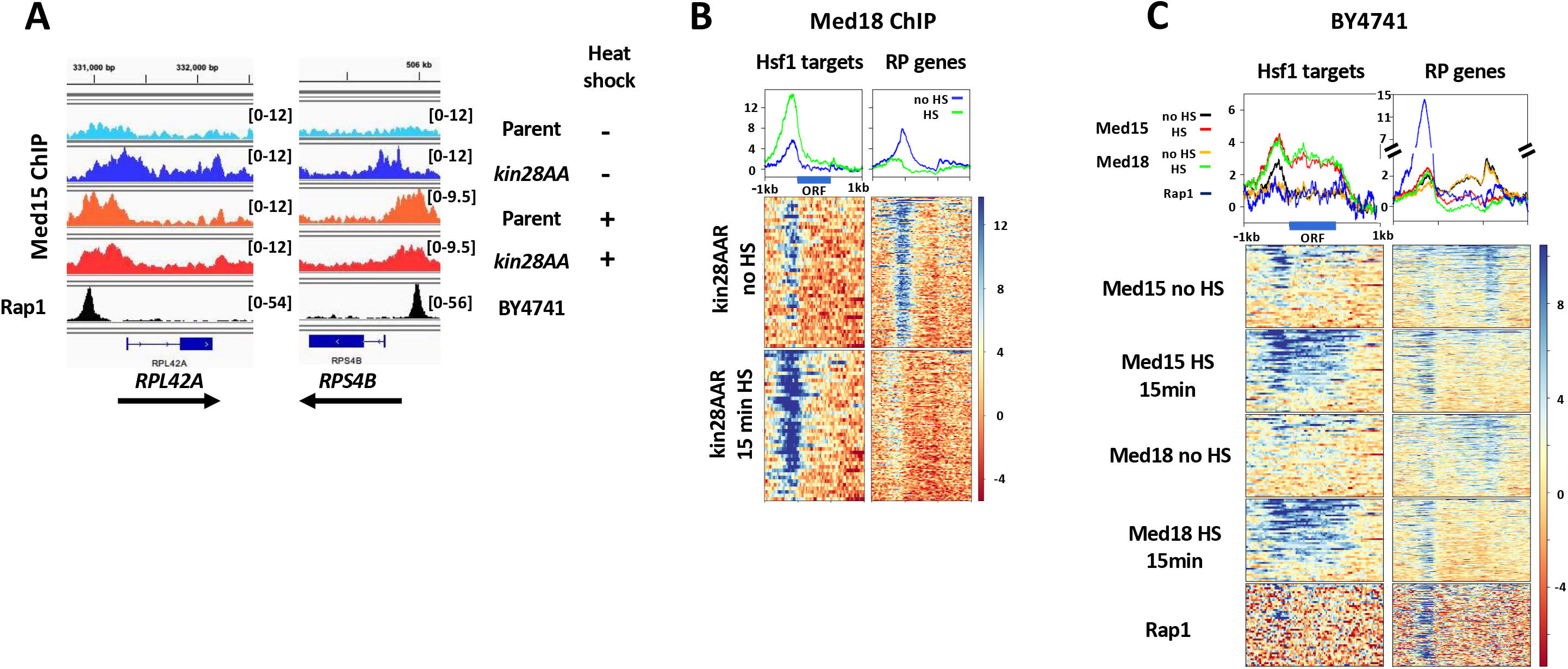
Mediator association persists at RP genes after heat shock. (A) Browser scans showing normalized occupancy of Med15 upstream of *RPL42A* and *RPS4B* in *kin28AA* yeast and the parent strain YFR1321, both treated with rapamycin, with and without heat shock (top four scans) or Rap1 in strain BY4741 (bottom scan). Scale, in reads per million mapped reads, is indicated for each scan. (B) Heat maps and line graphs depicting normalized occupancy of the Mediator head module subunit, Med18, in *kin28AA* yeast in the presence of rapamycin, with and without heat shock, as 42 Hsf1 targets and 137 RP genes. (C) Heat maps and line graphs depicting normalized occupancy of Mediator subunits Med15 (tail) and Med18 (head), and Rap1, at Hsf1 target genes and RP genes in BY4741. The baseline for the Rap1 line graph was rescaled to align with the other line graphs. The signal observed at ORF regions (seen at Hsf1 targets under heat shock conditions, and at RP genes under non-heat-shocked conditions) is a ChIP artifact frequently observed at highly transcribed ORFs (Eyboulet et al. 2013; Park et al. 2013; Teytelman et al. 2013; Jeronimo and Robert 2014; Knoll et al. 2020).

As mentioned earlier, the *kin28AA* strain and its parent, YFR1321, harbor the *tor1-1* mutation, which prevents the normal stress response accompanying rapamycin administration (Haruki et al. 2008). Because the Tor pathway is intimately connected with ribosome function (Mayer and Grummt 2006), we considered the possibility that the unexpected association of Mediator with RP genes repressed by heat shock was a consequence of the *tor1-1* mutation. We therefore performed ChIP-seq against Med15 and Med18 in the wild type strain BY4741 before and after 15 min of heat shock. Again, we observed peaks at RP genes for both Med15 and Med18 following heat shock (Figure 3C; Figure S3). Furthermore, as with the anchor-away parent strain YFR1321, Mediator ChIP-seq signal at RP genes was stronger after heat shock. The peaks for Med15 and Med18 were nearly coincident with the peak observed for Rap1, which binds upstream of 127 out of 137 RP genes and is essential for recruitment of the pre-initiation complex to RP genes (Figure 3C) (Lieb et al. 2001; Mencia et al. 2002; Ansari et al. 2009; Zeevi et al. 2011; Knight et al. 2014; Reja et al. 2015).

The ChIP-seq peaks for Mediator subunits observed upstream of RP genes in both wild type and *kin28-AA* yeast after heat shock suggest that Mediator is recruited to these genes in heat shocked cells but does not transit to proximal promoters as it normally does in non-heat-shocked cells. Many non-RP genes are also repressed by heat shock (Figure 1) (Gasch et al. 2000; Causton et al. 2001). To assess whether Mediator occupies promoters of such genes after heat shock, we examined ChIP-seq signal at UAS genes—that is, the 498 genes found to exhibit discernible Mediator occupancy in wild type yeast (Jeronimo et al. 2016). We divided these genes into those showing decreased Pol II occupancy by at least 2-fold (116 “UAS down” genes after removing RP genes) and the 231 “UAS not down” genes having a ratio of Pol II occupancy in heat shocked compared to non-heat-shocked cells ≥1, and compared the Med15 signal at these two sets of genes before and after heat shock (Figure 4A). We observed Med15 signal for both the “UAS down” and “UAS not down” cohorts in YFR1321 yeast (parent strain for *kin28AA*) grown at 30°C; upon heat shock, Med15 signal was unchanged for “UAS down” genes and showed a slight increase for “UAS not down” genes (Fig. 4A). Thus, these two cohorts behaved similarly with regard to Mediator occupancy in spite of their divergent behavior with regard to Pol II occupancy (Figure 4A).

**Figure 4.**
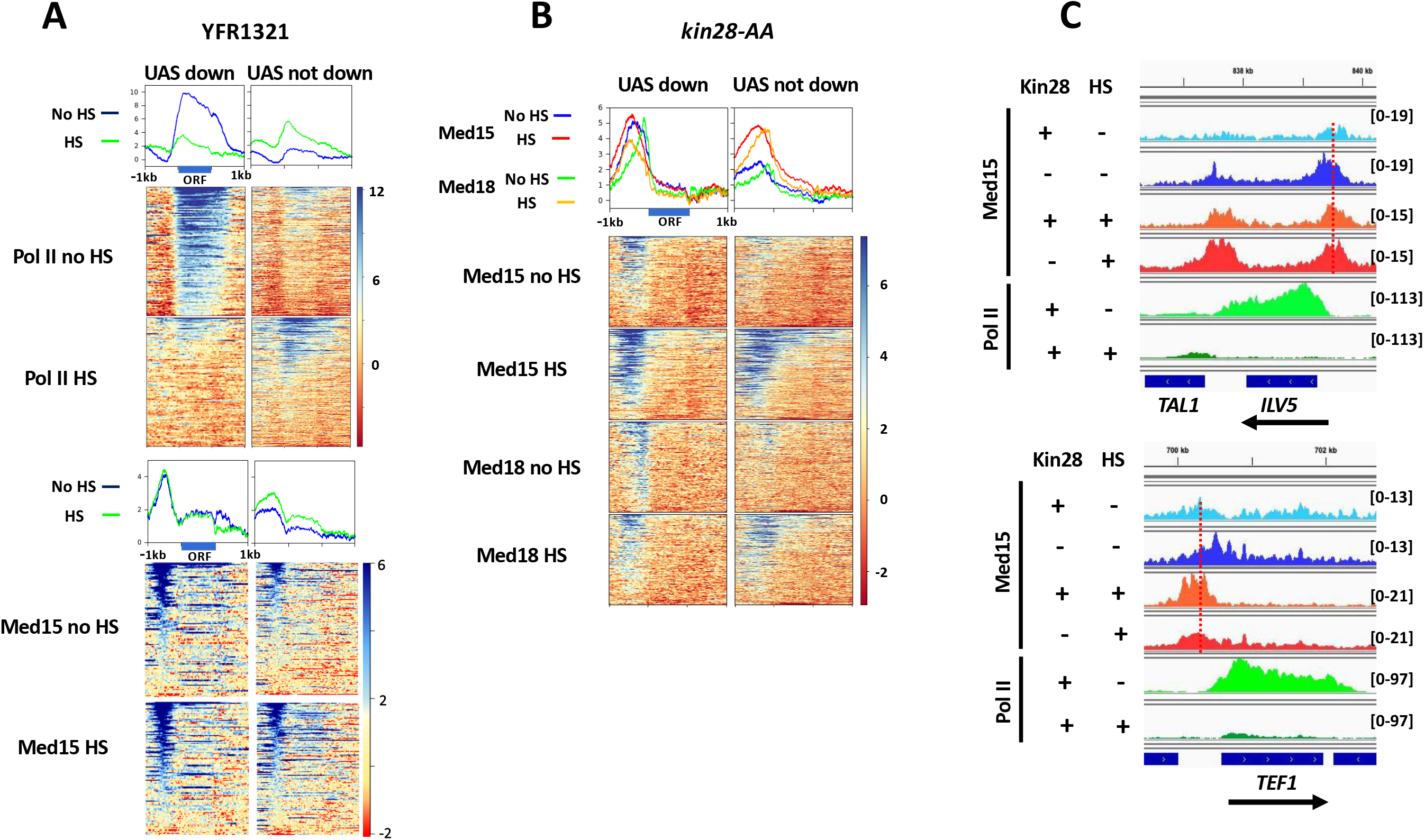
Mediator association at non-RP genes repressed by heat shock. (A) Heat maps and line graphs showing normalized occupany of Pol II and Med15 at UAS genes (see text) having Pol II occupancy decreased by at least 2-fold upon heat shock (“UAS down”) or having Pol II occupancy unchanged or increased upon heat shock (“UAS not down”) in YFR1321, the parent strain to *kin28-AA* yeast, with and without heat shock. (B) Heat maps and line graphs showing normalized occupany of Med15 (tail) and Med18 (head) at “UAS down” and “UAS not down” genes in *kin28-AA* yeast treated with rapamycin, with and without heat shock. (C) Browser scans showing Med15 and Pol II occupancy at *ILV1* and *TEF1* genes in *kin28-AA* yeast (“+ Kin28”) or the parent strain YFR1321 (“-Kin28”), both treated with rapamycin, with or without heat shock. Scale, in reads per million mapped reads, is indicated for each scan. Note that Pol II occupancy is reduced at both genes upon heat shock; the vertical dashed lines emphasize the shift of the Med15 peak towards the promoters only when Kin28 is depleted in the absence of heat shock.

We then examined association of Med15 and Med18 at these same cohorts under conditions of Kin28 depletion. Similar to our observations at RP genes, both subunits exhibited continued association with genes repressed by heat shock (Figure 4B, left panels, and Figure 4C). When cells were not heat shocked, Med18, from the head module, exhibited a narrow peak close to the TSS, while Med15, from the tail module, showed a broader peak extending from near the TSS to farther upstream. This is consistent with previous observations, and concordant with Mediator being recruited to upstream activating sites via the tail module and then transiting to the proximal promoter upon engagement of the head module with PIC components (Jeronimo et al. 2016; Petrenko et al. 2016; Knoll et al. 2018). Thus, because the tail module is in closer contact with the UAS and the head module in closer contact with the proximal promoter, cross-linking is more efficient for Med15 at the UAS and for Med18 near the TSS. Upon heat shock, both subunits exhibited peaks that were shifted upstream relative to the peaks seen in the absence of heat shock.

Altogether, these results indicate that Mediator remains associated with UAS regions of many genes that are repressed by heat shock, including RP genes, but is unable to recruit Pol II or to facilitate PIC formation (Vinayachandran et al. 2018).

### Role of PIC components in Mediator association following heat shock

We next addressed the role of PIC components TBP, TFIID, and Pol II in stabilizing Mediator occupancy at gene promoters in heat shocked cells. Previously we reported on the effect of simultaneous depletion of Taf1, TBP, or the Pol II subunit Rpb3 together with Kin28, compared to Kin28 alone, on association of Mediator with gene promoters in yeast growing in rich medium (Knoll et al. 2018). We examined the effects at two categories of gene promoters: TFIID-dominated genes, which lack consensus TATA elements, are relatively enriched for association with the Taf subunits of TFIID, and comprise ~85% of yeast genes, including most constitutively active genes; and SAGA-dominated genes, which possess consensus TATA elements, show relatively reduced Taf association, and are enriched for inducible genes (Huisinga and Pugh 2004; Tirosh and Barkai 2008; Rhee and Pugh 2012; Donczew et al. 2020; Knoll et al. 2020). Under conditions of Kin28 depletion, Mediator association was affected by deletion of any of the PIC components examined. Concordant with the distinction between TFIID-dominated and SAGA-dominated genes, depletion of Taf1 reduced association of Med15 and Med18 to a greater degree at TFIID-dominated genes than at SAGA-dominated genes, while Rpb3 depletion reduced Mediator association strongly at all genes (Knoll et al. 2018). In contrast, simultaneous depletion of TBP and Kin28, when compared to depletion of Kin28 alone, resulted in Mediator ChIP-seq peaks shifting upstream from the proximal promoter region to UAS sites at both SAGA-dominated and TFIID-dominated genes, indicating that TBP is required for transit of Mediator from its initial site of recruitment to its normally transient engagement with the proximal promoter. Similar effects are observed on comparing the effects of PIC depletion between TATA-containing, Taf1 depleted genes (enriched for SAGA-dominated genes) with TATA-less, Taf1 enriched genes (enriched for TFIID-dominated genes) (Rhee and Pugh 2012): Taf1 depletion reduces Mediator association at the latter class, and also at RP genes, much more than at TATA-containing, Taf1 enriched genes, while TBP depletion results in an upstream shift together with a decrease in intensity in the average ChIP-seq peaks for Med15 and Med18 for both categories (Figure 5A).

**Figure 5.**
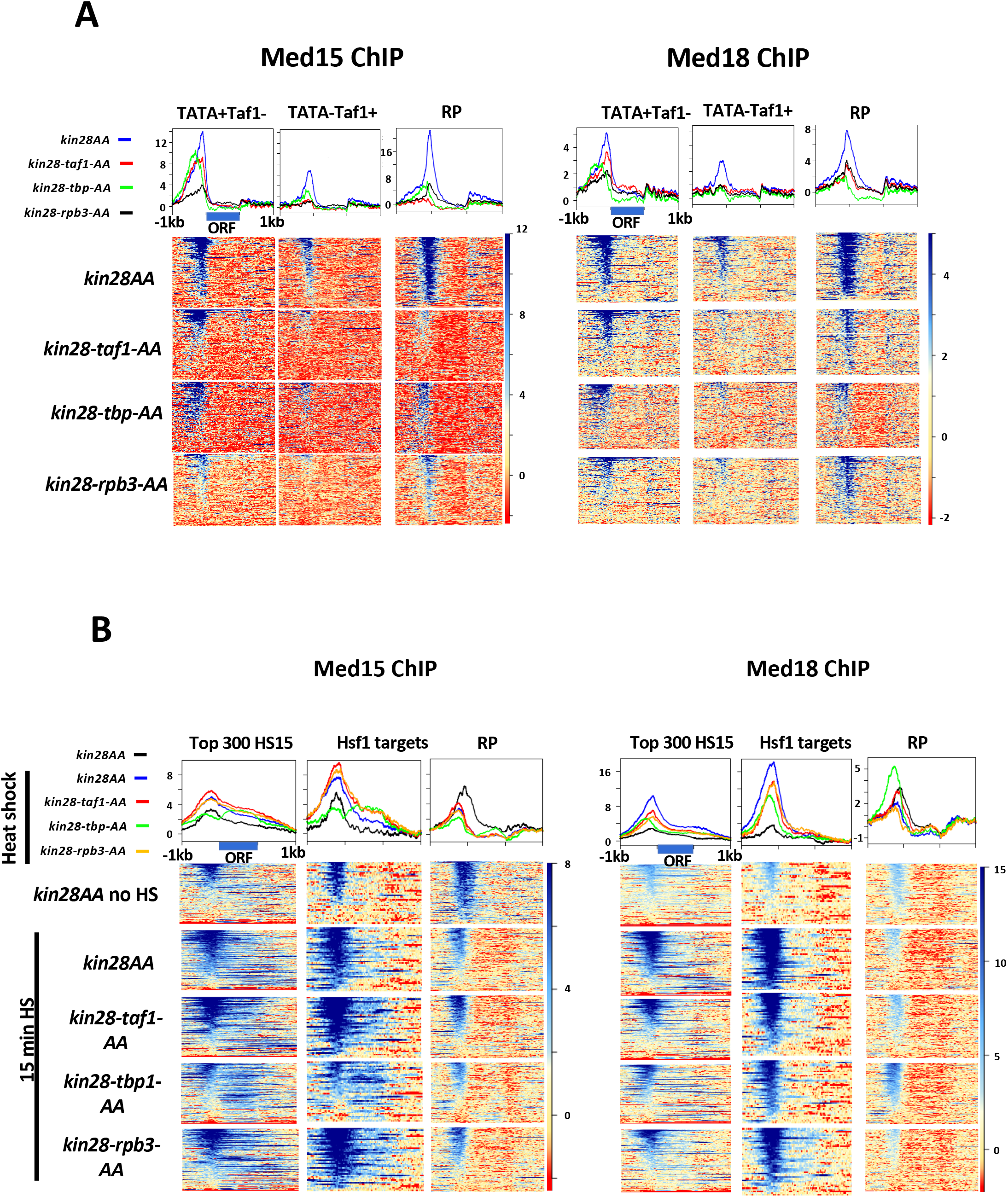
Effect of depleting PIC components on Mediator association with gene promoters. (A) Heat maps and line graphs showing normalized occupancy of Med15 (tail) and Med18 (head) at TATA-containing, Taf1 depleted promoters from the 1000 genes with highest Pol II occupancy (228 genes), TATA-less, Taf1 enriched genes excluding RP genes from the 1000 genes with highest Pol II occupancy (330 genes), and 137 RP genes after depletion of Kin28 alone or together with Taf1, TBP, or Rpb3, as indicated. (B) Heat maps and line graphs showing normalized occupancy of Med15 (tail) and Med18 (head) at the ~300 genes with highest Pol II occupancy after heat shock, 42 Hsf1 targets, and 137 RP genes after depletion of Kin28 alone or together with Taf1, TBP, or Rpb3, as indicated, without or with 15 min heat shock, as indicated.

To determine whether Mediator association is similarly dependent on PIC components in heat shocked cells, we performed ChIP-seq against Med15 and Med18 following heat shock in yeast cells depleted for Kin28 alone or in combination with Taf1, TBP, or Rpb3. Somewhat surprisingly, the ~300 genes showing highest Pol II occupancy after 15 min of heat shock were relatively insensitive to depletion of either Taf1 or Rpb3, in spite of comprising similar fractions of TATA-containing and TATA-less genes (111 and 83 genes, respectively) (Figure 5B). Indeed, the TATA-containing and TATA-less subsets from this cohort behaved nearly identically: both exhibited almost no change in Med15 ChIP-seq signal upon depletion of Taf1 or Rpb3, and modest decrease in Med18 signal (Figure S4). Depletion of TBP resulted in a slight upstream shift in both Med15 and Med18 ChIP-seq signals, consistent with the effect seen in the absence of heat shock (Figure 5B). (We note that ChIP-seq signal was seen over ORF regions in these experiments. This signal appears to be artifactual, although its origin is not completely understood (Eyboulet et al. 2013; Park et al. 2013; Teytelman et al. 2013; Jeronimo and Robert 2014; Paul et al. 2015b; Grunberg et al. 2016).

Hsf1 target genes similarly showed little effect of Taf1 or Rpb3 depletion on Med15 occupancy, while Med18 ChIP-seq signal showed a small decrease (Figure 5B). Depletion of TBP led to a similar decrease in Med18 signal along with a very slight shift upstream, and to a larger decrease in Med15 signal (Figure 5B). Hsf1 target genes have been noted to be nearly equally divided among TFIID-dominated and SAGA-dominated cohorts (de Jonge et al. 2017). Finally, Mediator signal at RP genes after heat shock appeared mostly insensitive to depletion of Taf1, TBP, or Rpb3, with the exception being an increased Med18 signal upon depletion of Kin28 together with TBP (Figure 5B). This insensitivity was not surprising, inasmuch as all three of these PIC components are depleted from RP genes under heat shock conditions (Figure 1) (Reja et al. 2015; Vinayachandran et al. 2018).

### Effect on Pol II and Mediator occupancy of exposure to cadmium

The unexpected observation of persistent Mediator association with genes repressed by heat shock prompted us to examine the effect of another environmental stress on association of Mediator with gene promoters. To this end, we chose to examine the effect of exposure to CdCl_2_. Exposure to this toxic metal at sub-millimolar concentrations induces a rapid transcriptional response in which ~150-500 genes are induced and from 18 to ~400 genes repressed (Momose and Iwahashi 2001; Cormier et al. 2010; Hosiner et al. 2014; Huang et al. 2016). Induced genes are enriched for binding sites for Hsf1, Msn2, and Msn4, and genes involved in ribosomal biogenesis are repressed by CdCl_2_ exposure, in common with the response to heat shock and other environmental stresses (Gasch et al. 2000; Causton et al. 2001; Hosiner et al. 2014). However, CdCl_2_ exposure also induces genes involved in sulfur compound metabolism (particularly the *MET* genes), among others, that are not involved in the heat shock response (Momose and Iwahashi 2001; Cormier et al. 2010; Hosiner et al. 2014; Huang et al. 2016), thus providing an environmental perturbation that results in a distinct but overlapping response to that elicited by heat shock.

We first tested the effect of CdCl_2_ exposure on Pol II association by ChIP-seq (Figure 6A). Marked induction of Pol II association was observed with the 47 genes showing the largest increase in mRNA abundance upon CdCl_2_ exposure (Momose and Iwahashi 2001), while association of Pol II with RP genes decreased to near baseline levels, in agreement with previous studies showing suppression of RP gene mRNA species upon CdCl_2_ exposure (Momose and Iwahashi 2001; Hosiner et al. 2014; Huang et al. 2016). Promoters showing at least 3-fold increase in normalized Pol II occupancy upon CdCl_2_ administration, and from the top 1000 Pol II-occupied genes after CdCl_2_ exposure, overlapped strongly with those identified in microarray studies (data not shown) (Momose and Iwahashi 2001). Gene ontology analysis revealed enrichment for categories related to sulfur compound metabolism and amino acid biosynthesis, and with response to stress, as expected (Table S2). Transcription factors enriched for binding to induced genes included Hsf1, Met4, Met32, Msn2, Msn4, and Yap1, all identified previously in a microarray study (Hosiner et al. 2014), as well as Atf2, Cbf1, Met31, and Skn7 (Table S2).

**Figure 6.**
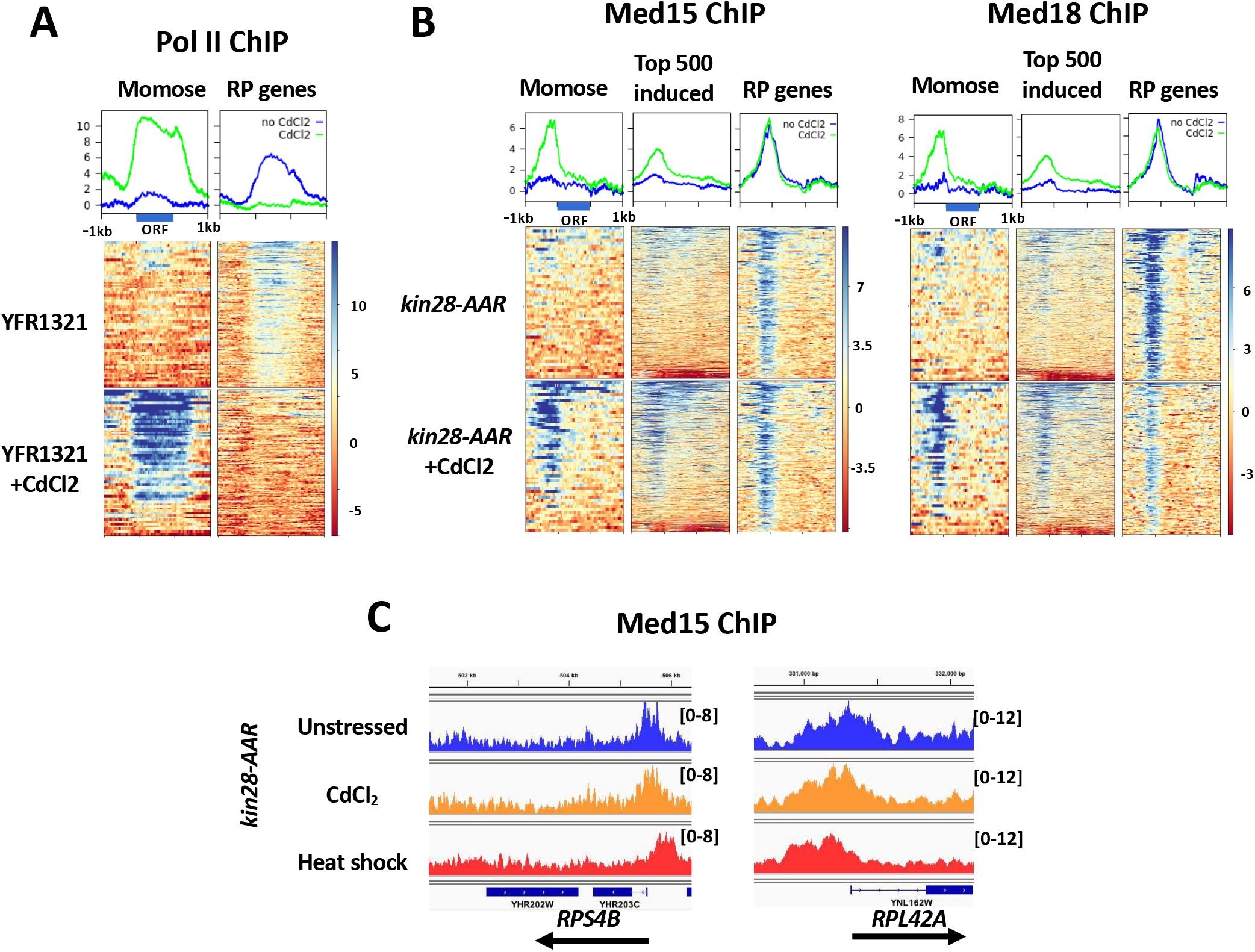
Effect on Pol II and Mediator occupancy of CdCl_2_ exposure. (A) Heat maps and line graphs showing normalized occupancy of Pol II at 47 strongly induced genes (Momose and Iwahashi 2001) and 137 RP genes in the anchor away parent strain, YFR1321. (B) Heat maps and line graphs showing normalized occupancy by Med15 (tail module) and Med18 (head module) at 47 strongly induced genes, the 500 genes having the highest ratio of induced to uninduced Pol II occupancy and 137 RP genes. (C) Browser scans showing Med15 occupancy in reads per million mapped reads after Kin28 depletion in unstressed cells, cells exposed to CdCl_2_, and after 15 min of heat shock, as indicated, at the *RPS4B* and *RPL42A* loci.

We next examined the effect of CdCl_2_ exposure on Mediator association. ChIP-seq of Med15, from the tail module, and Med18, from the head module, under conditions of Kin28 depletion, revealed increased association at genes identified as showing increased expression upon CdCl_2_ administration (Figure 6B, “Momose”) and at those showing increased Pol II association. Similarly to our observations of the effect of heat shock, RP genes, in spite of showing greatly reduced Pol II association upon CdCl_2_ exposure, exhibited persistent association of both Med15 and Med18 (Figure 6B). Also in accord with the effect of heat shock, Med18 exhibited relatively lower signal at RP genes after CdCl_2_ exposure than did Med15. Unlike Mediator association after heat shock, little if any upstream shift in the peaks for Med15 and Med18 at RP genes was observed after CdCl_2_ exposure (Figures 6B-C; compare Figures 2 and 3). To further compare heat shock and CdCl_2_ exposure, we examined the effect of CdCl_2_ exposure on Mediator and Pol II association at “UAS genes”, which we earlier defined as those identified as showing Mediator ChIP signal at UAS regions under conditions of active Kin28 (Jeronimo et al. 2016). We divided these genes into those showing decreased Pol II occupancy in the presence of CdCl_2_ and those having unchanged or increased occupancy (Figure S5). As with UAS genes following heat shock, Med15 and Med18 association persisted at genes showing decreased Pol II occupancy, while increasing (Med15) or staying constant (Med18) at genes having unchanged or increased Pol II occupancy (Figure S5). However, unlike the case for heat shock, and mirroring the results for RP genes, little or no shift of Mediator to more upstream sites was observed after CdCl_2_ exposure. We conclude that the stresses of heat shock and CdCl_2_ exposure both allow continued Mediator association with repressed genes while suppressing Pol II association, but that they differ in the extent to which they allow Mediator to remain associated with gene promoters rather than UAS regions under conditions of Kin28 depletion.

## Discussion

The Mediator complex was first identified as a coactivator needed to achieve activator-stimulated transcription in an *in vitro* system (Kornberg 2005). Consistent with this role, Mediator was shown to be recruited by activators (Kuras et al. 2003) and to facilitate looping between enhancers and promoters (Kagey et al. 2010). Recent work has added detail to this model, showing that in yeast, a single Mediator complex is first recruited to an upstream activation site and then, concomitant with its engagement with the PIC, transits to the proximal promoter region (Jeronimo et al. 2016; Petrenko et al. 2016). Association of Mediator with the proximal promoter is normally brief, but is stabilized by inhibiting Pol II escape either by inactivation or depletion of Kin28 or by loss of the Kin28 phosphorylation sites in the carboxy-terminal domain of Rpb1, the largest subunit of Pol II (Jeronimo and Robert 2014; Wong et al. 2014). Transit of Mediator from UAS to proximal promoter depends on TBP, and its association with the proximal promoter in the absence of Kin28 is destabilized by depletion of Taf1 or Pol II (Knoll et al. 2018).

Mediator occupancy of UAS regions of strongly induced genes, as assayed by ChIP, appears considerably stronger than at many constitutively active genes, even those expressed at high levels (Fan et al. 2006; Fan and Struhl 2009; Kim and Gross 2013). However, whether this reflects altered dynamics of Mediator with respect to the model described above has not been closely examined. In this work, we used ChIP-seq in combination with rapid depletion of Kin28 and PIC components to investigate Mediator association with activated and repressed genes following heat shock.

Consistent with previous work, we find increased association of Pol II and Mediator with genes induced by heat shock, with Mediator ChIP signal being evident even without Kin28 depletion (Figures 1-2) (Fan et al. 2006; Kim and Gross 2013; Petrenko et al. 2016; Petrenko et al. 2017). Pol II occupancy was reduced 2-3 fold upon depletion of the essential Med17 (head module) subunit of Mediator, both under both heat-shock and non-heat-shock conditions, and the reduction in occupancy did not differ for targets of Hsf1 or Msn2/4 as compared to other genes expressed under heat shock conditions (Figure 1C). Deletion of tail module subunits nearly abolishes induced Pol II occupancy at heat shock responsive genes (Kim and Gross 2013), while reduction in Pol II occupancy or nascent mRNA transcription of two-to eight-fold has been reported upon acute depletion or inactivation of essential Mediator subunits in yeast (Paul et al. 2015b; Plaschka et al. 2015; Petrenko et al. 2017; Warfield et al. 2017; Bruzzone et al. 2018). Thus, genes expressed during heat shock exhibit dependence on Mediator similar to genes expressed during normal growth. The relatively uniform dependence of gene expression on Mediator observed in yeast is not seen in mammalian cells, as depletion of core Mediator subunits most strongly affects a subset of genes, while little effect on nascent transcription is observed at many genes (Jaeger et al. 2020).

Depletion of Kin28, which inhibits promoter escape by Pol II (Wong et al. 2014), increases Mediator ChIP signal in the absence of heat shock; the effect is gene-dependent, with, for example, RP genes exhibiting a greater increase than do Hsf1 targets (Figure 2A). Under heat shock conditions, Med15 association is not increased at Hsf1 targets by Kin28 depletion, rather exhibiting a slight shift towards the promoter (Figure 2A-B and Figure S1). These observations suggest that Mediator dynamics may be altered during heat shock, such that its residence time and consequent “ChIP-ability” may increase relative to non-heat-shock conditions. Interestingly, a genetic screen uncovered mutations in several Mediator subunits that reduced the dynamic range in the transcriptional response to heat shock (Singh et al. 2006); whether these mutations might affect dynamics of Mediator is unknown.

An unexpected finding was the persistent association of Mediator with promoters of genes repressed, and having low or negligible Pol II association, following heat shock or exposure to CdCl_2_ (Figures 2, 3, 4, and 6). Mediator association with UASs of genes down-regulated by sulfometuron methyl treatment, which mimics amino acid starvation, was also observed in a study utilizing ChEC-seq, in which Mediator-tethered MNase is used to monitor association (Grunberg et al. 2016). Our results suggest that Mediator continues to be recruited to UASs of genes repressed by heat shock, but that recruitment of TBP and formation of the PIC is prevented by an unknown mechanism (Figure 7). The evidence in favor of this scenario is as follows: First, association of both Med15 from the tail module and Med18 from the head module is observed at RP genes and non-RP, “UAS genes”, that are repressed following heat shock, both in the *kin28-AA* parent strain YFR1321 as well as the lab strain BY4741 (Figures 2-4). In *kin28-AA* yeast treated with rapamycin, the observed ChIP-seq signals are shifted upstream relative to their positions in the absence of heat shock, and coincide closely with Rap1 binding sites at RP genes (Figure 3A and C); Rap1 is required for Mediator recruitment to most RP genes (Ansari et al. 2009). Second, the ChIP-seq signal for Med15 does not increase at repressed RP genes upon depletion of Kin28 (Figure 2A), and neither Med15 nor Med18 ChIP signals at UAS genes that are down-regulated by heat shock shift towards the proximal promoter upon depletion of Kin28 (Figure 4B). Pugh and colleagues have reported greatly reduced association of PIC components, including TBP, to genes repressed by heat shock (Vinayachandran et al. 2018). TBP appears to be required for transit of Mediator from its site of recruitment by activators to the proximal promoter (Knoll et al. 2018); thus, inhibition of PIC assembly by heat shock could prevent Mediator transit without preventing its initial recruitment, resulting in more stable association of Mediator with UAS regions at RP genes and other genes repressed by heat shock. Finally, the modest effect of depletion of TBP, Taf1, or Rpb3 on association of Med15 and Med18 with RP genes following heat shock is consistent with the lack of a PIC at these genes (Vinayachandran et al. 2018), and the engagement of Mediator at UAS regions where it would not interact with those components (Figure 5).

**Figure 7.**
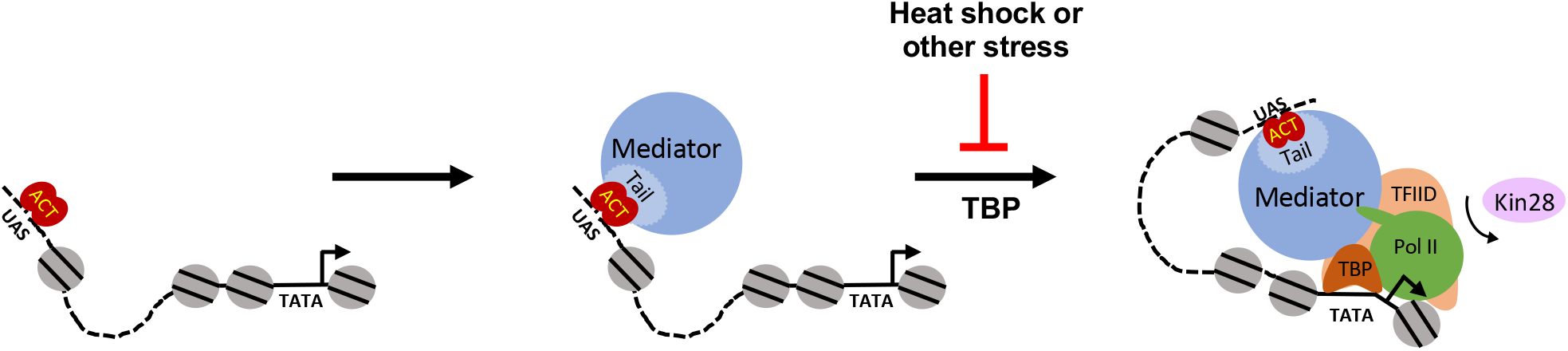
Cartoon of Mediator recruitment and transcriptional activation. Under normal growth conditions, Mediator is recruited via its tail module by an activator bound to a UAS, and in turn recruits components of the PIC, with TBP being required for Mediator transit from UAS to the proximal promoter, which may or may not possess a consensus TATA element. Association of Mediator with the proximal promoter is normally transient, being rapidly lost upon Kin28-dependent promoter escape by Pol II, but is stabilized by depletion or inactivation of Kin28. Heat shock or other stress (such as CdCl_2_ exposure) prevents PIC assembly and transit of Mediator to the proximal promoter through an unknown mechanism.

What mechanism allows Mediator association with repressed genes following heat shock while preventing PIC formation? Two possibilities seem most likely: 1) a factor or factors actively block assembly of the PIC, with consequent failure of Mediator to transit from the UAS to the proximal promoter (Knoll et al. 2018), or 2) activators or co-activators that are required for PIC assembly at heat-shock-repressed genes are inactivated by heat shock, while other factors (e.g. Rap1 at RP genes) remain that are able to recruit Mediator. In either case, the mechanism must allow transcription of those genes induced or not repressed by heat shock to be active, while repressing PIC assembly at a cohort of genes that includes, but is not limited to, RP genes.

At RP genes, Rap1 binds (at 129/137 RP genes) and recruits Fhl1 and Ifh1 to activate transcription (Martin et al. 2004; Schawalder et al. 2004; Wade et al. 2004; Rudra et al. 2005), while an additional factor, Sfp1, also associates with RP gene promoters in an Fhl1-dependent fashion (like Ifh1) and contributes to recruitment of TBP and Pol II (Jorgensen et al. 2004; Marion et al. 2004; Reja et al. 2015; Albert et al. 2019). In response to stress, a signaling cascade results in translocation of Crf1 into the nucleus, which competes with Ifh1 for binding to RP genes (Martin et al. 2004). Sfp1 is also evicted from RP genes and from the nucleus in response to stress, while Rap1 and Fhl1 remain associated with RP genes; the end result is inhibition of PIC assembly and concomitant loss of transcription (Jorgensen et al. 2004; Marion et al. 2004; Martin et al. 2004; Reja et al. 2015; Vinayachandran et al. 2018). Thus, Ifh1 and Sfp1 act as critical activators of RP gene expression. Rap1 and Fhl1 are required for recruitment of Ifh1 and Sfp1, but evidently are also able to recruit Mediator independently of Ifh1. This is surprising, as current models of transcriptional activation posit that Mediator recruitment leads directly to recruitment and assembly of the PIC. Further studies will be needed to understand the mechanism by which Mediator is recruited non-productively to genes repressed by heat shock or CdCl_2_, and to establish whether some “activators” may recruit Mediator but nonetheless be insufficient to facilitate PIC assembly and activate transcription.

The dynamics of Mediator recruitment and participation in PIC assembly have only recently begun to be appreciated, and thus far only in the model organism *Saccharomyces cerevisiae* (Jeronimo and Robert 2017). It is clear that in budding yeast, these dynamics vary in a gene-dependent fashion, as Mediator occupancy varies greatly at UAS regions and does not correlate with transcriptional output (Jeronimo and Robert 2014; Paul et al. 2015b; Grunberg et al. 2016). The work reported here shows that Mediator dynamics, including its association with repressed genes, also can vary in a condition-dependent fashion. Mediator itself is affected in its post-translational modifications and its composition by alterations in environment such as osmotic shock and the transition to stationary phase (Holstege et al. 1998; Miller et al. 2012); whether and how Mediator dynamics are affected by such alterations is currently obscure. Finally, in metazoan organisms, Mediator associates with enhancers that are sometimes many kilobases removed from the sites of PIC assembly at which Mediator participates via loop formation (Kagey et al. 2010); the dynamics of such Mediator-dependent loop formation, and the variables that influence those dynamics, remain an area for future investigation.

## Methods

### Yeast strains and growth

Yeast strains used in this study are listed in Table S3. Epitope tags were introduced into BY4741 to generate strains RMYDS1 and RMYDS2, and into YFR1321 to generate RMYDS10, by PCR amplification from strains containing the tagged protein and selectable marker, followed by transformation and selection (Hill et al. 1991; Longtine et al. 1998); strains were verified by PCR and ChIP. For simplicity, epitope-tagged strains are referred to by the parent strain names in the text and figures; for example, “BY4741” refers also to RMYDS1, which harbors the *med15-myc* allele. Cultures were grown in yeast peptone dextrose (YPD) medium (1% bacto-yeast extract, 2% bacto-peptone extract, 2% glucose). For Rap1 ChIP in wild type and *rap1-ts* yeast (Li et al. 2011), cultures were shifted to 37°C for 1 hr before cross-linking and ChIP and processed as described previously (Paul et al. 2015a). Anchor away experiments were carried out as reported previously (Knoll et al. 2018). In brief, yeast were grown to 0.6-0.8 OD_600_ at 30°C and then rapamycin (LC Laboratories, Woburn, MA, USA) was added from a 1 mg/ml stock solution in ethanol to a final concentration of 1μg/ml. One hour after rapamycin treatment, one-third volume of prewarmed media (13 ml at 57°C added to 39 ml culture for a final temperature of 37°C), or one-third volume of media at 30°C for non-heat-shocked cells, was added to each flask and flasks were immediately placed at 37°C (or at 30°C for non-heat-shocked cells) in a shaking incubator. Both *kin28-AA* yeast and the parent strain YFR1321 were treated with rapamycin in all experiments unless indicated otherwise. Samples were incubated for 15 min or 30 min prior to cross-linking with formaldehyde. Similarly, for CdCl_2_ treatment, yeast cultures were grown and treated with rapamycin as described above. After 1 hour of rapamycin treatment, CdCl_2_ was added from a 1 M stock solution to a final concentration of 0.5 mM and incubated for another hour before crosslinking.

### ChIP-seq

ChIP was performed as described previously (Knoll et al. 2018). For IP, 600 μL of WCE was incubated overnight at 4°C with 10μg of monoclonal RNA Poll antibody (Biolegend, USA), 2μg of anti-myc antibody (Sigma), or 2 μg of anti-Rap1 antibody (Santa Cruz, USA). Sixty microliters of WCE was used as Input control. Immunoprecipitated DNA was purified using 40 μL of protein A or G beads (Amersham/GE) with gentle agitation at 4°C for 90 min, and cleanup and purification performed as described previously (Knoll et al. 2018).

Libraries were prepared for sequencing using the NEBNext Ultra II library preparation kit (New England Biolabs, USA) according to manufacturer’s protocol and barcoded using NEXTflex barcodes (BIOO Scientific, Austin, TX, USA) or NEBNext Multiplex Oligos for Illumina. Purification and the size selection step were performed on barcoded libraries by isolating fragment sizes between 200 and 500 bp by using AMPureXP beads (Beckman Coulter, USA); size selection was confirmed by Bioanalyzer. Sequencing was performed at the Illumina NextSeq platform at the Wadsworth Center, New York State Department of Health (Albany, NY, USA) or at the Carolina Center for the Genome Sciences, University of North Carolina at Chapel Hill for Rap1 ChIP-seq. ChIP-seq experiments are summarized, and corresponding deposited files indicated, in Table S4. We also used previously published ChIP-seq data deposited at the NCBI Short Read Archive under accession number PRJNA413080 (Knoll et al. 2018).

### ChIP-seq analysis

Unfiltered sequencing reads were aligned to the *S. cerevisiae* reference genome (Saccer3) using bwa (Seoighe and Wolfe 1999). Up to 1 mismatch was allowed for each aligned read. Reads mapping to multiple sites were retained to allow evaluation of associations with non-unique sequences (Seoighe and Wolfe 1999) and paired end reads were removed, with the exception of the Rap1 ChIP-seq data, which yielded single-end reads and for which duplicate reads were retained. Calculation of coverage, as shown in heat maps and line graphs, was preceded by library size normalization, and was performed with the “chipseq” and “GenomicRanges” packages in BioConductor (Gentleman et al. 2004). Alternatively, reads were aligned and analysis conducted using the Galaxy platform (Goecks et al. 2010) and Excel. For metagene analysis, including heat maps, we subtracted reads from an input control (strain YFR1321, the parent strain to *kin28-*AA yeast (KHW127), grown in the absence of rapamycin); for Rap1, reads from the *rap1-2 ts* mutant, which has greatly reduced binding at 37°C (Drazinic et al. 1996; Ganapathi et al. 2011), were subtracted from the wild type strain also grown at 37°C for one hour. Pol II occupancy was determined as read depth over ORFs, and Mediator occupancy was determined as normalized read depth over the 300 bp upstream of TSS using BedCov in SamTools (Li et al. 2009). Clustering analysis (Figure 1) was performed using Cluster and Treeview (Eisen et al. 1998). The 1000 genes having highest Pol II occupancy, normalized to gene length, were obtained using BedCov to obtain read depth over coding sequences using Pol II ChIP-seq data (strain BY4741 grown at 30°C in YPD medium) (Paul et al. 2015b; Knoll et al. 2018). Genes designated as SAGA-dominated and TFIID-dominated were obtained from (Huisinga and Pugh 2004), and genes designated as containing or not containing a consensus TATA element, and being Taf1-enriched or Taf1-depleted, were obtained from (Rhee and Pugh 2012). Targets of Hsf1 were defined as those genes that are not activated by heat shock if Hsf1 is depleted (Pincus et al. 2018; Tye et al. 2019), Msn2/4 targets were defined in (Solis et al. 2016), and CdCl2 induced genes were those identified in Table 3 of (Momose and Iwahashi 2001). Occupancy profiles were normalized for read depth and generated using the Integrative Genomics Viewer (Robinson et al. 2011). Gene ontology analysis was performed using the Generic Gene Ontology Term Finder (https://go.princeton.edu/cgi-bin/GOTermFinder/GOTermFinder) (Boyle et al. 2004). Hypergeometric test p-values were calculated using the online calculator at http://www.alewand.de/stattab/tabdiske.htm.

## Supporting information

Table S3-Yeast strains

Table S4-ChIP-seq summary

## Data access

ChIP-seq reads have been deposited in the NCBI Short Read Archive (https://www.ncbi.nlm.nih.gov/sra) under project number PRJNA657372.

## Acknowledgements

We thank Francois Robert and Kevin Struhl for generously providing yeast strains, Jason Lieb and Colin Lickwar for help in ChIP-seq of Rap1, and Elisabeth Knoll for helpful discussions. We gratefully acknowledge help from the Wadsworth Center Applied Genomics Technology and Tissue Culture and Media Cores. This work was supported by the National Science Foundation (MCB1516839 to RHM) and in part by the NIH Intramural Research Program at the National Library of Medicine (ZIZ and DL).

## Disclosure declaration

No conflicts of interest declared.

## Notes

### Competing Interest Statement

The authors have declared no competing interest.

