## Supplementary Figures for "Altered Mediator dynamics during heat shock in budding yeast"

#### Supplementary Figure Legends

**Figure S1.** Effect of heat shock on Mediator association. (A) Heat maps and line graphs depicting normalized occupancy of the Mediator tail module subunit, Med15, in *kin28AA* yeast treated with rapamycin ("kin28AAR") and the parent strain YFR1321, also treated with rapamycin, before and after 15 min heat shock, at 42 Hsf1 targets and 213 Msn2-4 targets (see Methods) and 137 RP genes. (B) Box and whisker plots showing the ratios of Med15 occupancy with and without Kin28 depletion for the ~300 genes showing highest Med15 occupancy in Kin28-depleted cells without or with heat shock; ratios are also shown for RP genes (137 genes), and for Hsf1 targets (42 genes) and Msn2-4 targets (213 genes) in heat-shocked cells. The p-value for comparison of the ratios for the top 300 genes with and without heat shock was calculated using the Wilcoxon rank sum test. (C) Box and whisker plots showing the ratios of Med15 occupancy with and without Kin28 depletion for the ~300 genes showing highest occupancy by Pol II or Med15, as indicated, without or with heat shock. Data same as used for Figure 1. (D) Line graphs depicting occupancy of Med15 in kin28-depleted yeast (TBY128), normalized to the parent strain RMY1321-G, in each of two replicate experiments without and with heat shock. Occupancy was determined for the ~300 genes most highly occupied by Pol II after heat shock, and the 641 most highly occupied genes in the absence of heat shock; the latter number was chosen based on having the same lower cutoff for occupancy as for the heat shock cohort.

**Figure S2.** Browser scans showing Med15 occupancy in *kin28AA* yeast and the parent strain RMY1321-G, both treated with rapamycin, with and without heat shock. Scale, in reads per million mapped reads, is indicated for each scan. (A) *TPS1* and *TSL1* are Msn2/4 targets and are expressed at low levels in the absence of heat shock and induced 5-10 fold upon heat shock based on Pol II occupancy; (B) *TSA1* is expressed in the absence of heat shock and shows 2-fold increased Pol II occupancy on heat shock, and is not a target of Hsf1 or Msn2/4; and (C) *YDJ1* is a target of Hsf1 and is expressed both with and without heat shock with little change in Pol II occupancy. Note that *RPS18B* and *RPS17A* in (B) are strongly repressed upon heat shock.

**Figure S3.** Comparison of Mediator occupancy in epitope-tagged strains derived from wild type yeast (BY4741; strains RMYDS1 and RMYDS2) and after Kin28 depletion (kin28AAR; strains TBY128 and EKY18), with and without heat shock, at 42 Hsf1 targets, 213 Msn2-4 targets, and 137 RP genes. Left, Med15 (tail module) occupancy; right, Med18 (head module) occupancy.

**Figure S4.** Effect of depleting PIC components on Mediator association with gene promoters of TATA-containing and TATA-less promoters. Heat maps and line graphs showing normalized occupancy of Med15 (tail) and Med18 (head) at TATA-containing promoters (111 genes) and TATA-less promoters (83 genes) from the 300 genes with highest Pol II occupancy after 15 min of heat shock. Occupancy is shown after depletion of Kin28 alone with or without heat shock, and after depletion of Kin28 and Taf1, TBP, or Rpb3 followed by 15 min of heat shock.

**Figure S5.** Effect on Pol II and Mediator occupancy of $CdCl_2$ exposure at UAS genes. Heat maps and line graphs showing normalized occupancy of Pol II at 498 genes identified as exhibiting Mediator ChIP-seq peaks in wild type yeast (Jeronimo et al., 2016), separated into genes showing Pol II occupancy reduced by at least 2-fold and excluding RP genes ("Down", 40 genes) or having Pol II occupancy between 80% and 125% seen in uninduced cells ("Not down", 286 genes). Pol II occupancy was determined in the anchor away parent strain YFR1321 with or without $CdCl_2$ exposure, and Med15 (tail module) and Med18 (head module) occupancies were measured in *kin28-AA* yeast treated with rapamycin with or without $CdCl_2$ exposure.

Jeronimo, C., Langelier, M.F., Bataille, A.R., Pascal, J.M., Pugh, B.F., and Robert, F. (2016). Tail and Kinase Modules Differently Regulate Core Mediator Recruitment and Function In Vivo. Mol Cell *64*, 455-466.

#### Figure S1

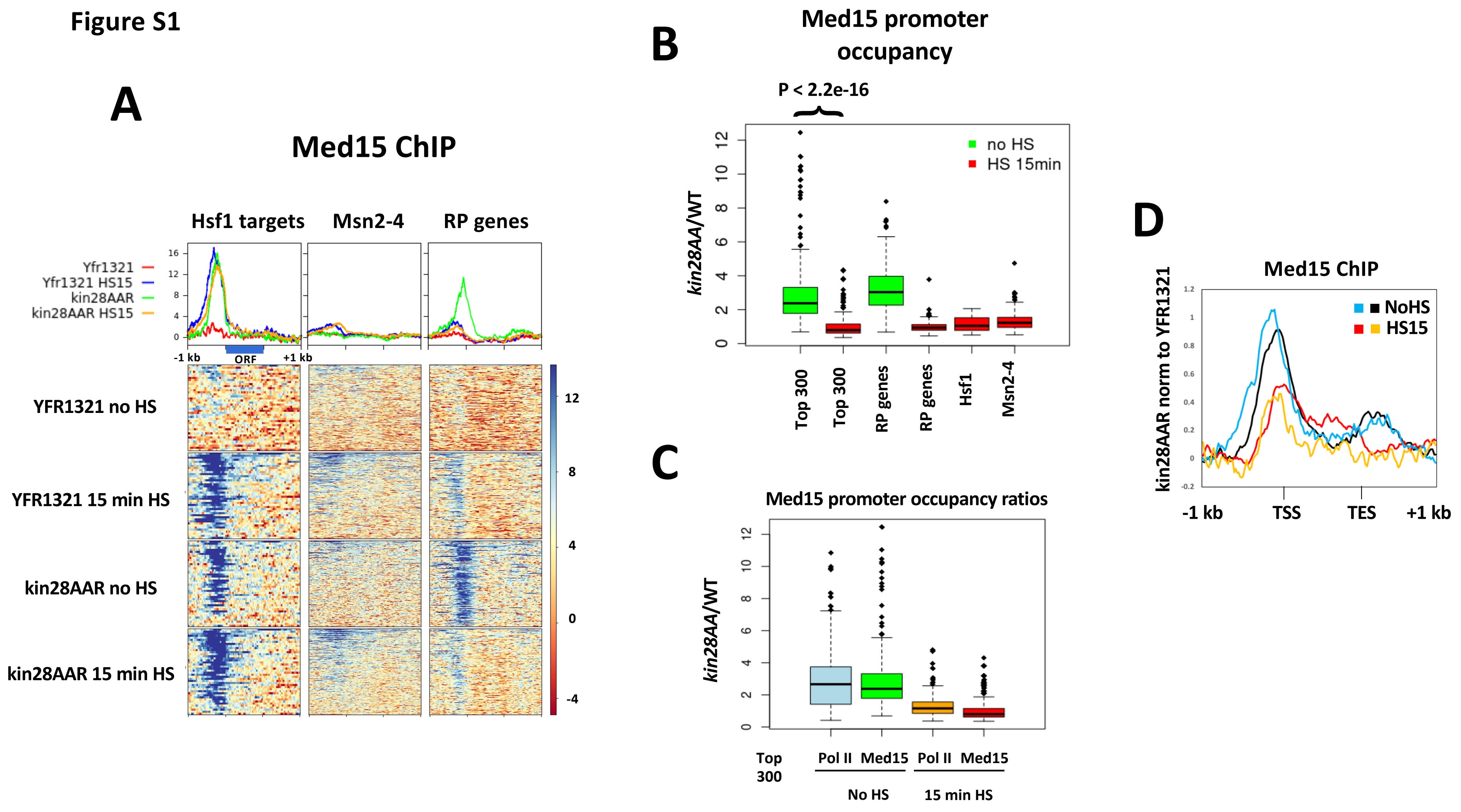

A
Med15 ChIP
Hsf1 targets
Msn2-4
RP genes
Yfr1321
Yfr1321 HS15
kin28AAR
kin28AAR HS15
-1 kb
ORF
+1 kb
YFR1321 no HS
YFR1321 15 min HS
kin28AAR no HS
kin28AAR 15 min HS
B
Med15 promoter occupancy
P < 2.2e-16
kin28AA/WT
no HS
HS 15min
Top 300
Top 300
RP genes
RP genes
Hsf1
Msn2-4
C
Med15 promoter occupancy ratios
kin28AA/WT
Top 300
Pol II
Med15
Pol II
Med15
No HS
15 min HS
D
Med15 ChIP
kin28AAR norm to YFR1321
NoHS
HS15
-1 kb
TSS
TES
+1 kb

#### Figure S2

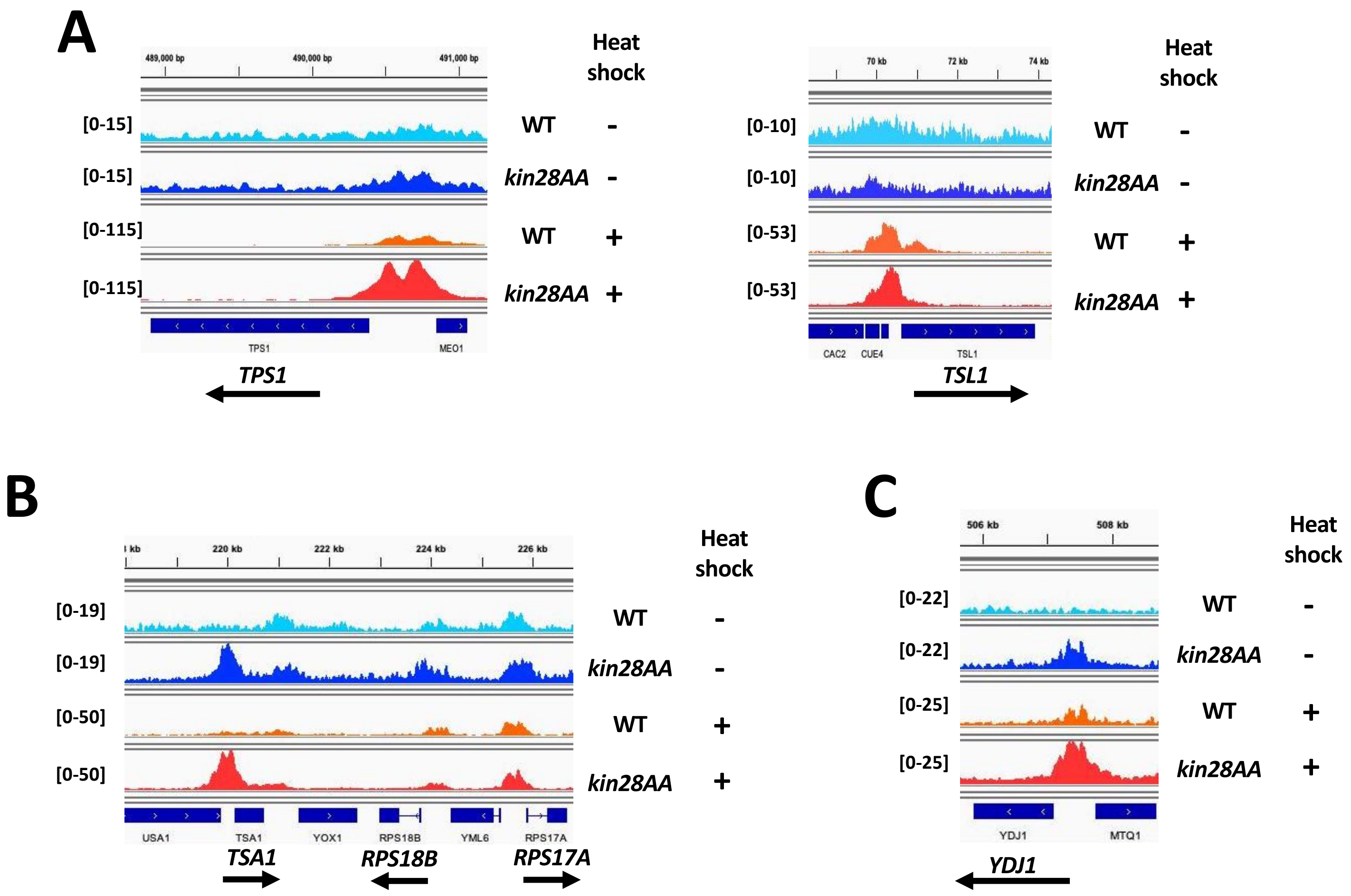

A
489,000 bp
490,000 bp
491,000 bp
Heat shock
[0-15]
WT -
[0-15]
kin28AA -
[0-115]
WT +
[0-115]
kin28AA +
TPS1
MEO1
TPS1
70 kb
72 kb
74 kb
Heat shock
[0-10]
WT -
[0-10]
kin28AA -
[0-53]
WT +
[0-53]
kin28AA +
CAC2
CUE4
TSL1
TSL1
B
220 kb
222 kb
224 kb
226 kb
Heat shock
[0-19]
WT -
[0-19]
kin28AA -
[0-50]
WT +
[0-50]
kin28AA +
USA1
TSA1
YOX1
RPS18B
YML6
RPS17A
TSA1
RPS18B
RPS17A
C
506 kb
508 kb
Heat shock
[0-22]
WT -
[0-22]
kin28AA -
[0-25]
WT +
[0-25]
kin28AA +
YDJ1
MTQ1
YDJ1

### Figure S3

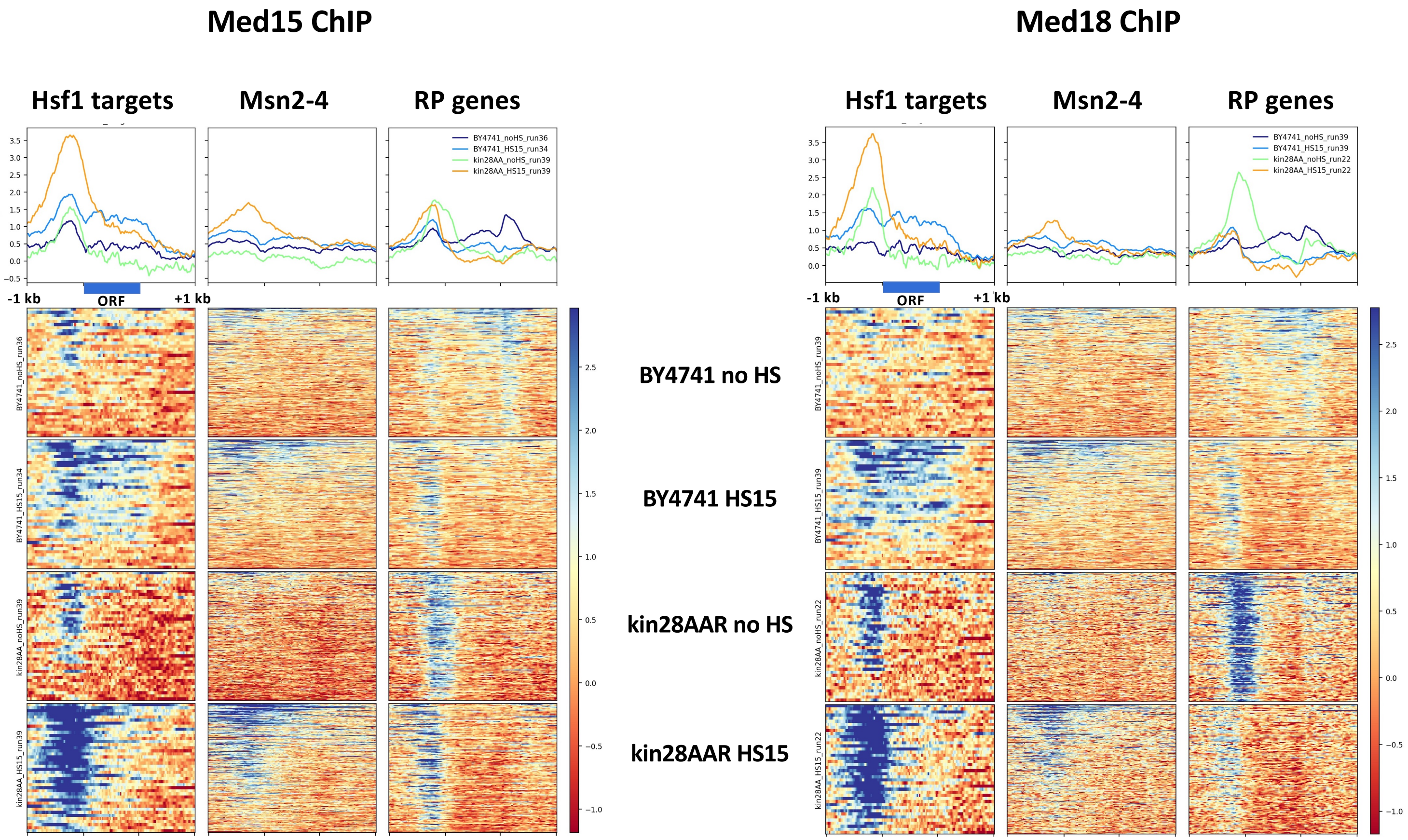

Med15 ChIP
Med18 ChIP
Hsf1 targets
Msn2-4
RP genes
BY4741_noHS_run36
BY4741_HS15_run34
kin28AA_noHS_run39
kin28AA_HS15_run39
BY4741_noHS_run39
BY4741_HS15_run39
kin28AA_noHS_run22
kin28AA_HS15_run22
-1 kb
ORF
+1 kb
BY4741 no HS
BY4741 HS15
kin28AAR no HS
kin28AAR HS15

#### Figure S4

##### Top 300 Pol II occupied genes after 15 min HS

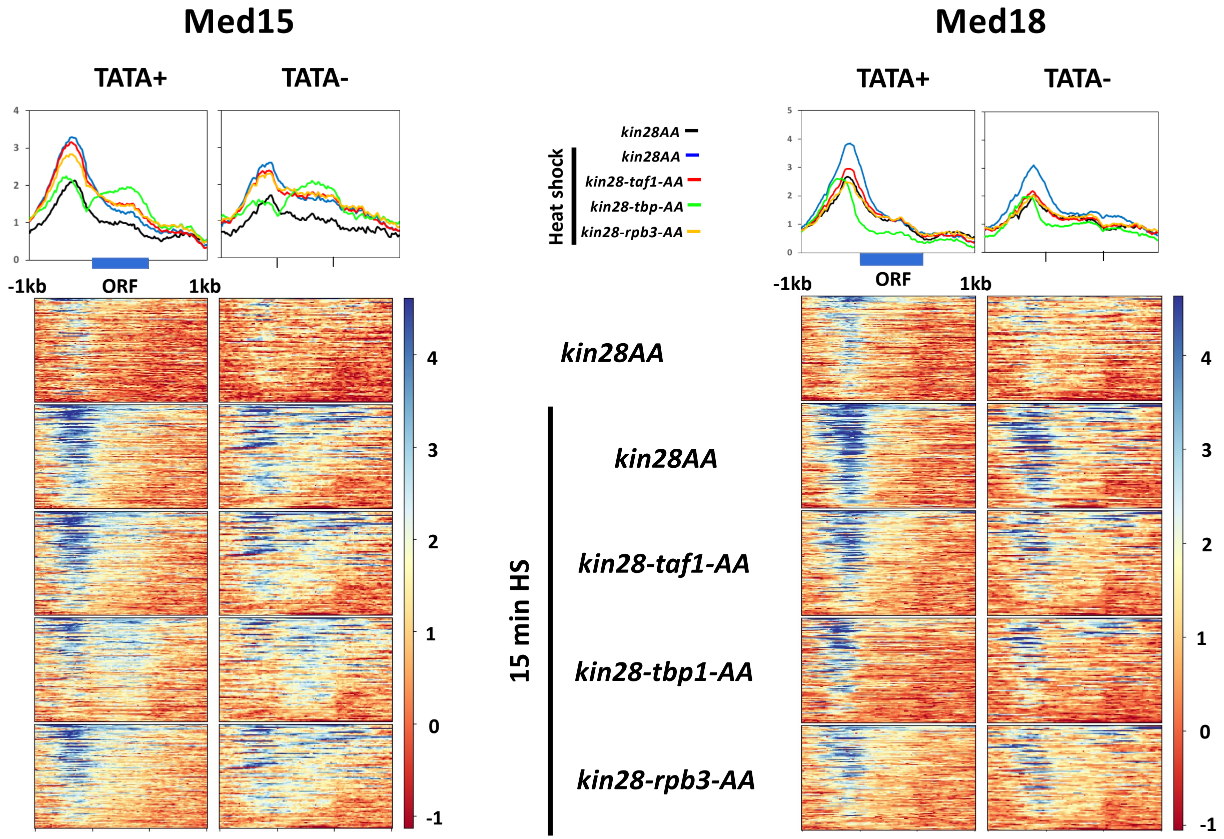

### Figure S5

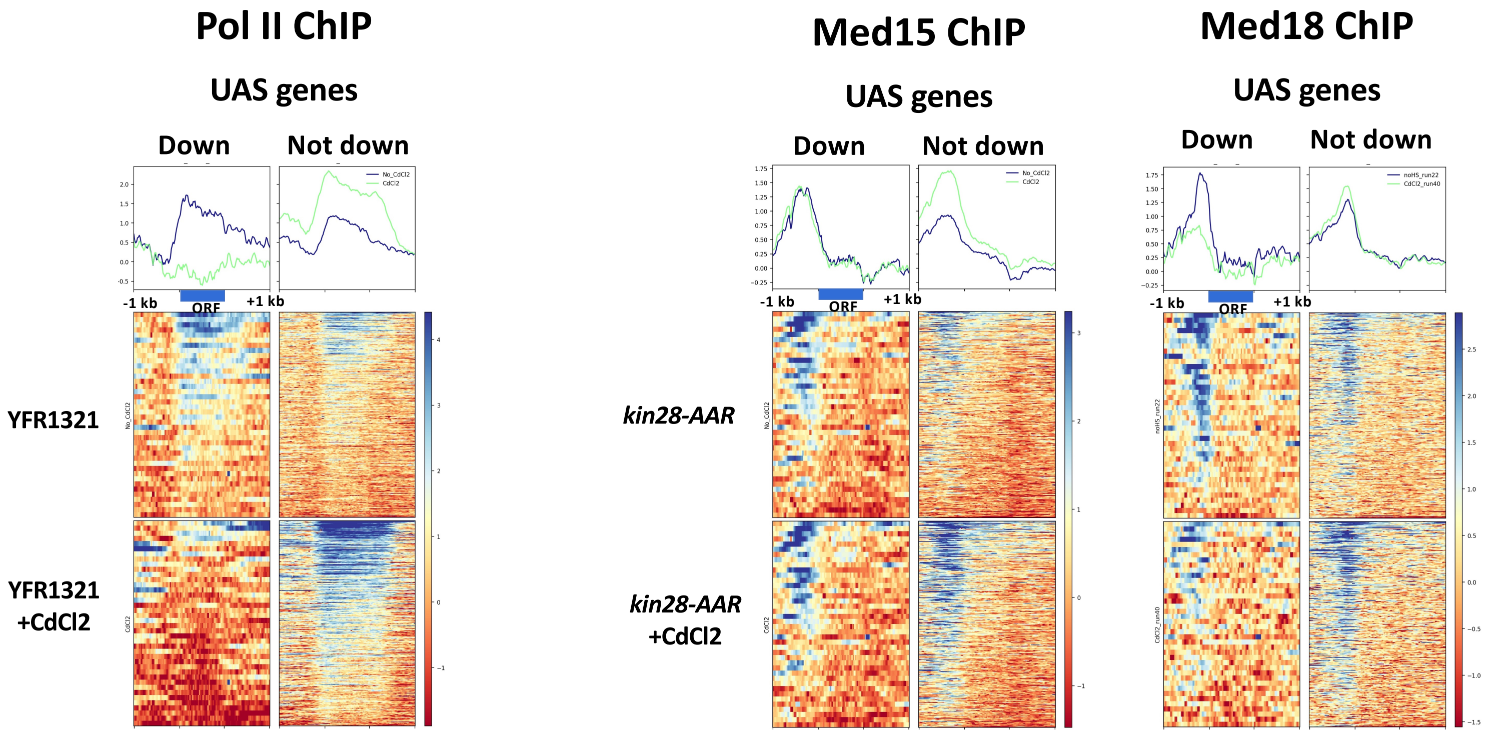

Pol II ChIP
UAS genes
Down
Not down
-1 kb
ORF
+1 kb
YFR1321
YFR1321
+CdCl2
Med15 ChIP
UAS genes
Down
Not down
-1 kb
ORF
+1 kb
kin28-AAR
kin28-AAR
+CdCl2
Med18 ChIP
UAS genes
Down
Not down
-1 kb
ORF
+1 kb
